# Transposon insertion sequencing identifies genetic determinants of intrinsic rifamycin resistance in *Mycobacterium abscessus*

**DOI:** 10.64898/2026.06.10.731268

**Authors:** Kotaro Sawai, Hidetada Hirakawa, Takehiko Mima, Yuta Morishige, Yasuhiro Maeda, Motoko Shinohara, Satoshi Mitarai, Yohei Doi, Yusuke Minato

## Abstract

*Mycobacterium abscessus* exhibits intrinsic resistance to many antimicrobial agents, including rifampicin, a frontline anti-tuberculosis drug, severely limiting treatment options. Here, we used transposon insertion sequencing (Tn-Seq) to perform a genomewide screen to identify genes required for intrinsic rifampicin resistance in *M. abscessus*. We uncovered candidate genes that confer intrinsic resistance to the rifamycin-class antibiotics, rifampicin and rifabutin. The genes we identified included previously reported genes such as *arr*, *helR,* and *MAB_2807*. By comparing our results with the rifampicin intrinsic resistance gene in *Mycobacterium tuberculosis*, we found that the mechanisms underlying rifampicin intrinsic resistance were distinct between the two species. The contribution of seven representative candidate genes to rifampicin resistance was confirmed by characterizing targeted gene deletion or transposon insertion mutants. Among these determinants, *MAB_2807* was identified as a major efflux-based contributor to rifamycin resistance, and its disruption increased intracellular rifampicin accumulation. In addition to known resistance determinants, Tn-Seq revealed contributions from many genes involved in cell envelope processes to rifamycin resistance. Guided by this genetic signature, we evaluated the combined effects of cell wall-targeting antimicrobial agents with rifamycins. We found that rifamycins displayed selective synergistic interactions with specific β-lactam antibiotics. Interestingly, we found that rifamycin exposure altered cell envelope ultrastructure and increased the accumulation of the peptidoglycan precursor UDP-N-acetylglucosamine, suggesting that rifampicin perturbs envelope-associated metabolic homeostasis. These findings highlight rifamycin-induced vulnerable cellular processes that may inform rational combination strategies.

**Importance:** Rifamycins are cornerstone antibiotics for the treatment of mycobacterial infections, yet they are ineffective against *Mycobacterium abscessus*. Although several key resistance mechanisms have been described, the full genetic basis of intrinsic rifamycin resistance remains to be elucidated. Here, we used transposon insertion sequencing to systematically identify genes that contribute to intrinsic rifamycin resistance in *M. abscessus*. Because recent studies have shown that chemical modification of rifamycins can overcome specific intrinsic resistance mechanisms, a comprehensive understanding of these determinants could facilitate the development of next-generation rifamycins with improved activity against *M. abscessus*. In addition, we unexpectedly found that rifamycin exposure alters cell envelope structure, the intracellular levels of a key metabolic intermediate in peptidoglycan biosynthesis, and susceptibility to certain β-lactams in *M. abscessus*. These findings may help guide the rational development of combination therapeutic strategies with next-generation rifamycins.

## Introduction

*Mycobacterium abscessus* is an emerging opportunistic pathogen that causes chronic pulmonary and soft tissue infections, with a rapidly increasing global incidence over the past two decades (1–3). Among nontuberculous mycobacteria, *M. abscessus* is recognized as one of the most drug-resistant species and poses a substantial therapeutic challenge. Its intrinsic resistance necessitates prolonged multidrug regimens and is associated with high rates of treatment failure and relapse (4–6).

Rifampicin is a cornerstone antimicrobial agent for the treatment of tuberculosis, but is ineffective against *M. abscessus* infections. Rifabutin, a related rifamycin, has demonstrated modestly greater activity than rifampicin against *M. abscessus* (7); however, it remains less effective compared to its activity against *Mycobacterium tuberculosis,* the causative agent of tuberculosis. Rifamycins inhibit bacterial transcription by binding to the β-subunit of RNA polymerase (RpoB). In *M. tuberculosis*, resistance is primarily mediated by mutations within the 81-bp rifampicin resistance-determining region (RRDR) of *rpoB* (8). In contrast, *M. abscessus* exhibits high-level intrinsic resistance in the absence of *rpoB* RRDR mutations. To date, at least five genes, *arr*, *helR*, *MAB_2807*, *MAB_1915,* and *MAB_2362,* have been reported to confer rifampicin resistance in *M. abscessus* (9–14). Arr is the ADP-ribosyltransferase that inactivates rifampicin by ADP-ribosylation at the C25 position (11). HelR has been proposed to function as an RNA polymerase protective factor that counteracts rifampicin-mediated transcriptional inhibition. *MAB_2807* is a major facilitator superfamily (MFS) transporter that was shown to confer resistance to rifampicin, rifabutin, tetracycline, vancomycin, clofazimine, and ofloxacin (14). In addition, *MAB_1915* and *MAB_2362* have been associated with altered rifampicin and rifabutin susceptibility, although they are generally considered broader intrinsic resistance determinants rather than rifamycin-specific resistance factors (12, 13).

Recent studies have demonstrated that chemical modification of rifamycins can overcome specific intrinsic resistance mechanisms. For example, rifabutin derivatives that evade Arr-mediated ADP-ribosylation have demonstrated enhanced antibacterial activity against *M. abscessus* (15, 16). These findings highlight the potential of rationally designed rifamycin derivatives to circumvent defined resistance pathways and further develop them for the treatment of *M. abscessus* infection. However, these mechanisms have largely been investigated individually, and the genomewide architecture that enables survival under rifamycin stress has not been systematically defined. A comprehensive understanding of these determinants may facilitate the development of next-generation rifamycins with improved activity against *M. abscessus*, as well as combination strategies that potentiate rifamycin efficacy.

Here, we applied genomewide transposon insertion sequencing (Tn-Seq) under subinhibitory rifamycin stress to define the genetic determinants of intrinsic rifamycin resistance in *M. abscessus*. This approach identified a high-confidence core set of intrinsic rifamycin resistance genes that were required for both rifampicin and rifabutin resistance. We revealed that not only the canonical drug-modifying enzymes and RNA polymerase-associated factors, but also envelope-associated and physiological maintenance genes confer intrinsic rifamycin resistance in *M. abscessus*. Furthermore, we show that rifamycin exposure induces reproducible alterations in envelope structure that could account for selective synergistic interactions with specific β-lactam antibiotics. Collectively, our findings provide a systematic foundation for understanding how rifamycin stress reshapes bacterial physiology and identify cellular pathways that may be selectively exploited in rational combination therapy.

## Results

### Genomewide identification of intrinsic rifampicin and rifabutin resistance genes in *M. abscessus*

We generated a saturated transposon insertion mutant library of *M. abscessus* subsp. *abscessus* JCM 13569^T^ (*Mab* ATCC 19977) by collecting more than 300,000 individual transposon insertion mutants. Genomic DNA was extracted from each library, and transposon-adjacent regions were enriched by PCR prior to Illumina sequencing. Sequencing data were analyzed using the TRANSIT software package. The insertion saturation rates of biological replicates ranged from 46.4% to 56.5%, exceeding the recommended threshold for robust Tn-Seq analysis (i.e., >30% of TA sites disrupted) (Table S1). Gene essentiality was first assessed using the Hidden Markov Model (HMM) implemented in TRANSIT. Among 4,348 chromosomal genes analyzed, we identified 388 essential genes (ES), 173 growth advantage genes (GA), 134 growth defect genes (GD), and 3,653 non-essential genes (NE). The overall distribution of essentiality categories was consistent with previous Tn-Seq studies in *M. abscessus* (Table S1) (17, 18). On a plasmid pMAB23, nearly all genes were classified as non-essential, with the exception of three growth-defect genes (Table S2).

To identify genes that confer intrinsic resistance to rifampicin and rifabutin, we first observed growth of the *Mab* ATCC 19977 strain on 7H9 agar plates containing various concentrations of rifampicin and rifabutin (Figure S1). We then grew the *Mab* ATCC 19977 transposon insertion mutant library on 7H9 agar plates containing 5 µg/ml rifampicin, 0.5 µg/ml rifampicin, or 0.5 µg/ml rifabutin, concentrations that did not inhibit the growth of the *Mab* ATCC 19977 (Figure 1A, Figure S1). We used the two rifampicin concentrations to capture genetic requirements across different levels of rifampicin stress. In parallel, rifabutin was used to identify determinants shared across rifamycin-class antibiotics. The Tn-Seq data were analyzed using the resampling method implemented in TRANSIT to statistically test differences in insertion abundance between a drug-treated sample and its untreated control. Insertion saturation rates under all drug-treated conditions were comparable to those of the untreated control libraries (Table S2). We identified 91 intrinsic resistance candidate genes in the presence of 5 µg/ml rifampicin (Fig. 1B and Table S3). Of note, no resistance-associated genes were detected on the plasmid. Among the 91 genes, 28 genes were identified in the presence of 0.5 µg/ml rifabutin and 16 genes were identified in the presence of 0.5 µg/ml rifampicin (Fig. 1B and Table S4-5). Previously reported rifamycin resistance genes in *Mab* ATCC 19977, *arr*, *helR, MAB_2362*, and *MAB_2807*, were included in the 16 genes, suggesting that our screening was comprehensive (Table 1) (9–14).

**Figure 1.**
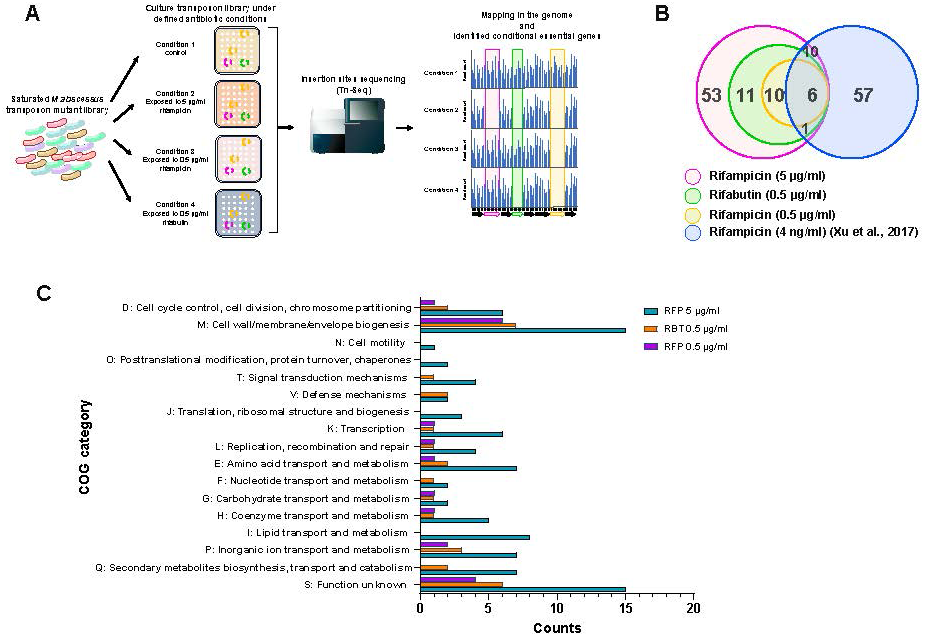
Genomewide identification of intrinsic rifamycin resistance genes by Tn-Seq. A) Experimental workflow to identify *M. abscessus* genes required for survival under rifamycin stress and intrinsic resistance. B) Venn diagram showing the number of genes identified as intrinsic rifamycin resistance determinants under each condition (adjusted *p* value < 0.05). C) Distribution of identified genes across COG functional categories.

**Table 1.** Functional annotation of the core set of 16 genes commonly required for intrinsic rifamycin resistance.

| RefSeq locus tag | genbank locus tag | gene | gene description |
| --- | --- | --- | --- |
| *MAB_RS01040* | *MAB_0177* | - | alpha/beta hydrolase |
| *MAB_RS02885* | *MAB_0539* | - | DUF6779 domain-containing protein |
| *MAB_RS03145* | *MAB_0591* | *arr* | NAD(+)-rifampin ADP-ribosyltransferase |
| *MAB_RS05485* | *MAB_1055c* | - | M23 family metallopeptidase |
| *MAB_RS05655* | *MAB_1087* | *glp* | gephyrin-like molybdotransferase Glp |
| *MAB_RS07890* | *MAB_1530* | *ldtMab2* | L,D-transpeptidase |
| *MAB_RS10105* | *MAB_1974* | *ripC* | peptidoglycan hydrolase RipC |
| *MAB_RS12045* | *MAB_2362* | *steA* | putative cytokinetic ring protein SteA |
| *MAB_RS12065* | *MAB_2366* | *xerD* | site-specific tyrosine recombinase XerD |
| *MAB_RS12280* | *MAB_2408c* | - | substrate-binding domain-containing protein |
| *MAB_RS13845* | *MAB_2728c* | - | NlpC/P60 family protein |
| *MAB_RS14240* | *MAB_2807* | - | MFS transporter |
| *MAB_RS16075* | *MAB_3170c* | - | M50 family metallopeptidase |
| *MAB_RS16170* | *MAB_3189c* | *helR* | RNA polymerase recycling motor ATPase HelR |
| *MAB_RS17895* | *MAB_3526c* | - | DUF3152 domain-containing protein |
| *MAB_RS20420* | *MAB_4026c* | - | TetR/AcrR family transcriptional regulator |

We examined their functional composition using gene classification based on the Clusters of Orthologous Groups (COG) functional categories. This analysis revealed a strong enrichment of genes associated with cell wall/membrane/envelope biogenesis (COG category M), which constituted the largest functional group across all three conditions (Figure 1C). In addition to envelope-related functions, a subset of genes was assigned to categories involved in cell cycle control, cell division and chromosome partitioning (COG category D) and transcriptional regulation (COG category K), indicating that intrinsic rifamycin resistance is supported by multiple physiological processes.

### Identification of *M. abscessus*-specific intrinsic rifampicin resistance genes

Xu *et al*. previously performed a similar Tn-Seq study to identify intrinsic resistance genes in *M. tuberculosis* (19). To assess the conservation of rifampicin resistance mechanisms between *M. tuberculosis* and *M. abscessus*, we compared our Tn-Seq dataset with the previously reported *M. tuberculosis* H37Rv dataset by Xu *et al*. Overall, only a limited overlap was observed between the two species. Among the 16 core intrinsic resistance genes identified in *Mab* ATCC 19977, only 6 genes were conserved in *M. tuberculosis* H37Rv, including *MAB_1087 (glp)*, *MAB_1974 (ripC)*, *MAB_2408c*, *MAB_2807*, *MAB_3170c*, and *MAB_3526c* (Figure 1B, Table S6). In contrast, most intrinsic resistance determinants were species-specific. We identified 10 genes unique to *Mab* ATCC 19977, including *arr* and *helR*, both of which lack orthologs in *M. tuberculosis*. Conversely, a large number of intrinsic resistance genes identified in H37Rv (n = 57) were not detected in our datasets. These include genes involved in lipid metabolism, cell envelope biosynthesis, and transport systems, such as *fecB* (*Rv3044*) and components of phosphate and lipid transport pathways.

### Functional characterization of *M. abscessus* intrinsic rifampicin resistance genes

In addition to the Tn-Seq analysis, we performed transposon insertion mutant screening and identified three *Mab* ATCC 19977 rifampicin-susceptible transposon insertion mutants. The mutants had transposon insertions in *MAB_2408*, *MAB_2807*, and *MAB_3526*. The identified three genes were included in the 16 genes that confer intrinsic resistance in all conditions (Fig. 1B and Table 1). Additionally, we selected 4 genes, *arr*, *helR*, *ldtMab2* (*MAB_RS07890*; *MAB_1530*), and *ripC* (*MAB_RS10105*; *MAB_1974*) from the 16 genes and constructed gene deletion mutants in *Mab* ATCC 19977. For targeted gene deletion, we used the ORBIT system (20), but our initial attempt failed. We suspected the tetracycline-inducible promoter did not function sufficiently in *Mab* ATCC 19977 and thus replaced it with an acetamide-inducible promoter. Using this modified vector, we successfully constructed gene-deletion mutant strains. All mutant strains exhibited increased susceptibility to both rifampicin and rifabutin compared with the wild-type strain, although the extent of sensitization varied among the genes (Table 2).

**Table 2.** Comparisons of rifamycin MICs with Tn-Seq fitness defect of each gene defecteve mutant strain.

| Strains | Tn-Seq fitness defect* | Rifampicin<br>MIC (µg/ml) | Rifabutin<br>MIC (µg/ml) |
| --- | --- | --- | --- |
| *Mab* ATCC 19977 (WT) | 0 | 32 | 4 |
| WT *MAB_2408::Tn* | -3.17 | 8 | 2 |
| WT *MAB_2807::Tn* | -4.23 | 4 | 1 |
| WT *MAB_3526::Tn* | -2.66 | 12 | 1.5 |
| WT *Δarr* | -7.2 | 0.25 | 0.016 |
| WT *ΔldtMab2* | -1.63 | 16 | 1 |
| WT *ΔripC* | -3.01 | 4 | 1 |
| WT *ΔhelR* | -3.1 | 1 | 0.125 |

\* Log<sub>2</sub> fold change at rifampicin (0.5 µg/ml)

**Table 3.**
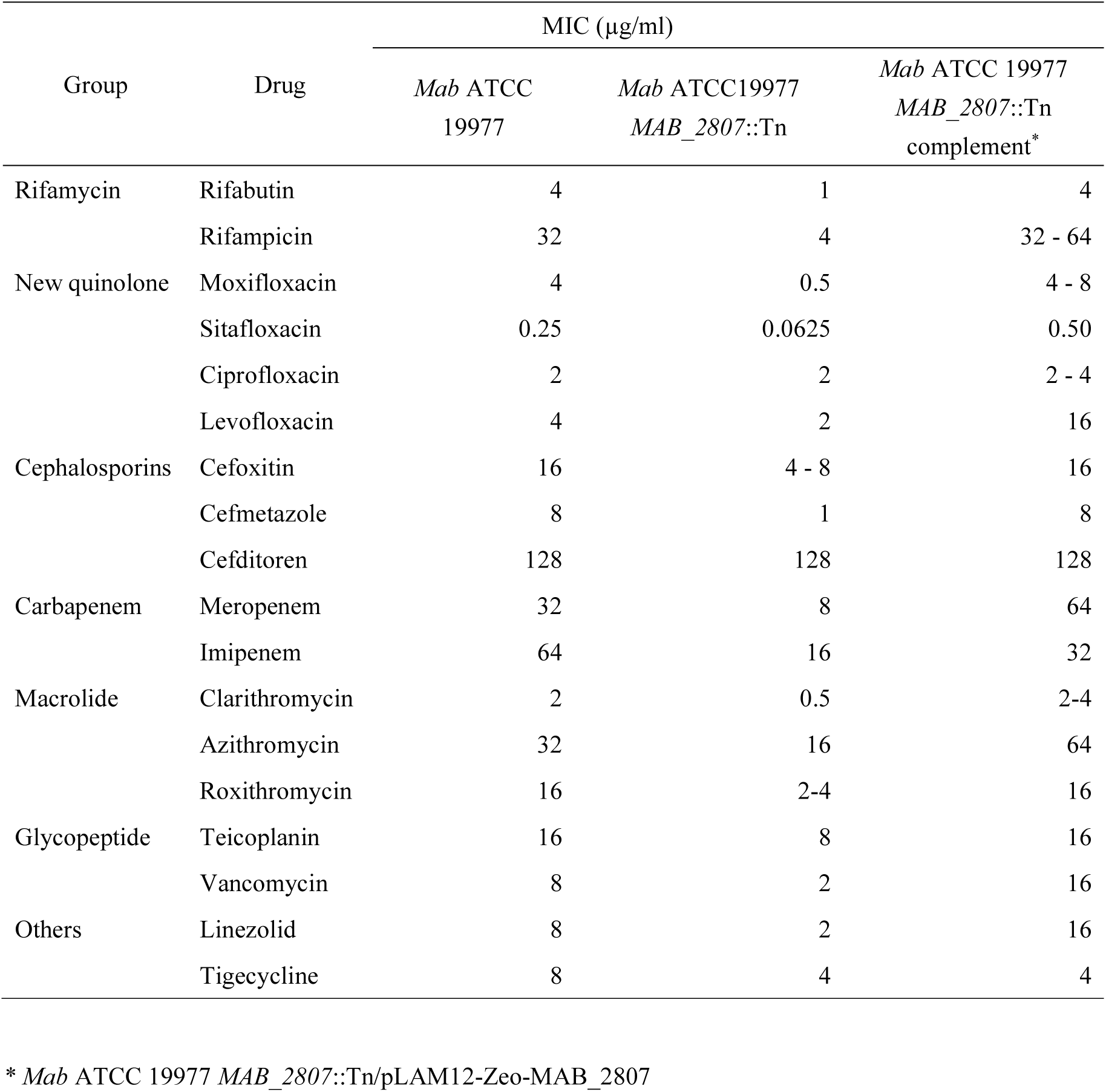
MICs of *Mab* ATCC 19977 and *MAB_2807*::Tn mutant strains.

To quantitatively evaluate the correlation of the Tn-Seq data, we examined the relationship between Tn-Seq fitness defects and growth fitness of individual mutants under the same drug concentrations used in the Tn-Seq experiments (Figure 2A). Across all three rifamycin selection conditions, the log₂ fold changes calculated by TRANSIT showed significant correlations with growth fitness, with adjusted R² values exceeding 0.5 and p-values < 0.03. We further assessed the relationship between Tn-Seq fitness defects and rifampicin and rifabutin minimum inhibitory concentration (MIC) values (Figure 2B). These correlations were strongest under lower drug concentrations, i.e. rifampicin (0.5 µg/ml) and rifabutin (0.5 µg/ml) with adjusted R² values exceeding 0.6 with *p*-values < 0.02. Collectively, these results demonstrate that Tn-Seq fitness scores quantitatively predict rifamycin susceptibility and validate the robustness of the core set of intrinsic resistance genes.

**Figure 2.**
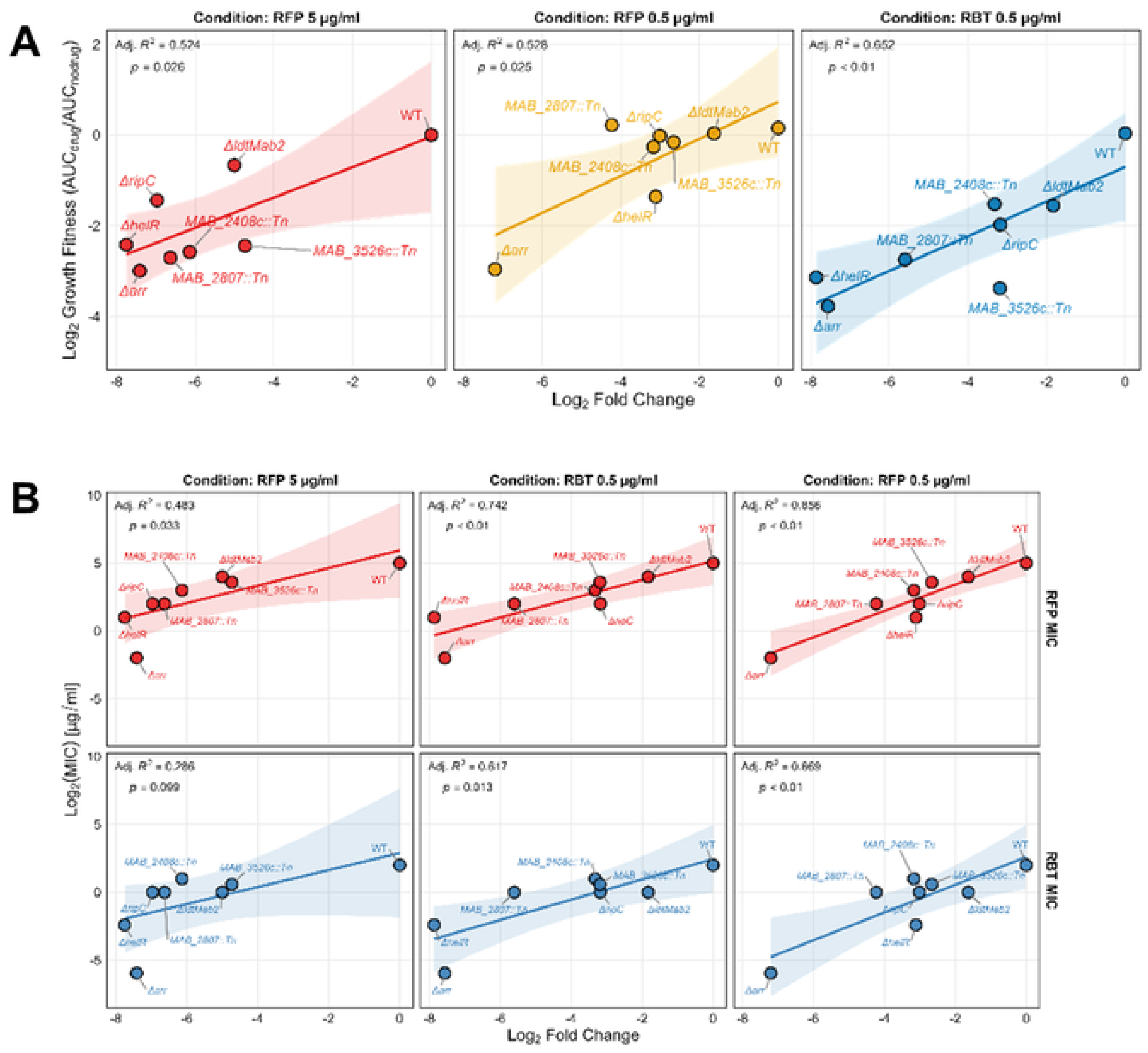
Quantitative correlation between Tn-Seq fitness defects and growth fitness under rifamycin stress or rifamycin MICs. A) Panels show the correlation between log₂ fold change values derived from Tn-Seq and experimentally measured growth fitness under three rifamycin conditions. Each point represents an individual mutant, plotted as the log₂-transformed growth fitness against the corresponding Tn-Seq log₂ fold-change value under the indicated condition. B) Upper panels show the correlation between rifampicin MICs and log2 fold change values derived from Tn-Seq under three rifamycin conditions analyzed using TRANSIT. Lower panels show the correlation between rifabutin MICs and log2 fold change values derived from Tn-Seq under the same conditions. Each point represents the log2-transformed MIC of an individual mutant plotted against its corresponding log2 fold change value under the indicated condition.

*MAB_2807*, an MFS transporter, was previously shown to confer multidrug resistance in *M. abscessus*, with reported effects on susceptibility to tetracycline, vancomycin, rifampicin, rifabutin, clofazimine, novobiocin, and ofloxacin (14). To extend these observations, we performed comprehensive drug susceptibility testing across a broader panel of antimicrobial agents, including fluoroquinolones (moxifloxacin, sitafloxacin, ciprofloxacin, and levofloxacin), β-lactams (cefoxitin, cefmetazole, cefditoren, meropenem, and imipenem), macrolides (clarithromycin, azithromycin, and roxithromycin), as well as teicoplanin, linezolid, and tigecycline. Consistent with previous reports, the *MAB_2807*::Tn mutant exhibited increased susceptibility to rifampicin and rifabutin. Notably, the mutant also displayed increased susceptibility to multiple additional antimicrobial agents spanning diverse classes that have not been previously evaluated in the context of *MAB_2807*, with fold increases ranging from 2- to 8-fold compared with the wild-type strain (Table 2). Among these drugs, rifamycins showed the largest fold changes in MIC. Genetic complementation of *MAB_2807* fully restored drug resistance in the *MAB_2807*::Tn mutant, confirming that the observed phenotypes were specifically attributable to the disruption of this gene. Furthermore, to directly assess whether *MAB_2807* contributes to rifampicin resistance by mediating drug efflux, we performed rifampicin accumulation assays using the wild-type strain and the *MAB_2807*::Tn mutant. Compared with the wild-type control, the *MAB_2807*::Tn mutant exhibited a significant increase in intracellular rifampicin accumulation (Figure S2). These results demonstrate that *MAB_2807* functions as an efflux transporter that reduces intracellular rifampicin levels, thereby contributing to rifampicin resistance. Together with the broad-spectrum drug susceptibility observed in the mutant, these data establish *MAB_2807* as a key efflux-based determinant of intrinsic rifamycin resistance in *M. abscessus*.

### Cell envelope-associated resistance signatures predict selective synergy with β-lactams

Given the strong enrichment of cell envelope-associated genes among intrinsic rifamycin resistance determinants, we evaluated the combinational effects of cell wall-targeting antimicrobial agents with rifampicin by using checkerboard assays (Figure 3, Figure S3). Drug interactions were assessed by the fractional inhibitory concentration index (FICI). As expected from our Tn-Seq results, rifampicin MICs decreased at least 4-fold when combined with cell wall-targeting antimicrobial agents, whereas those of cell wall-targeting antimicrobial agents decreased only 2-fold. Interestingly, MICs of certain β-lactams, imipenem (IPM) and cefditoren (CDTR), decreased at least 4-fold when combined with rifampicin. As a result, IPM and CDTR exhibited synergistic interactions with rifampicin (FICI ≤ 0.5), whereas other cell wall inhibitors showed additive interactions with rifampicin in *Mab* ATCC 19977. A similar trend was observed when the same drug combinations were tested against *M. abscessus* subsp. *abscessus* JCM 6390 strain. We also evaluated the combined effects of cell wall-targeting antimicrobial agents with fidaxomicin, another RNA polymerase inhibitor (21). We found that fidaxomicin also exhibited synergistic interactions with IPM and CDTR and additive interactions with the other cell-wall inhibitors tested. To confirm whether inhibition of RNA polymerase increases *M. abscessus* susceptibility to IPM and CDTR, we constructed a CRISPRi *rpoB* knockdown strain of *Mab* ATCC 19977 and measured MICs of cell wall targeting antimicrobial agents (Fig. S3). The *rpoB* knockdown strains showed increased susceptibility to IPM and CDTR, indicating that RNA polymerase inhibition is sufficient to confer selective vulnerability to these β-lactams (Figure S2). Notably, the rifamycin-β-lactam synergy was maintained in *arr*, *helR*, MAB_2807::Tn, and *ldtMab2* mutant backgrounds, indicating that this phenomenon is not driven by individual resistance determinants (Table S8, S9).

**Figure 3.**
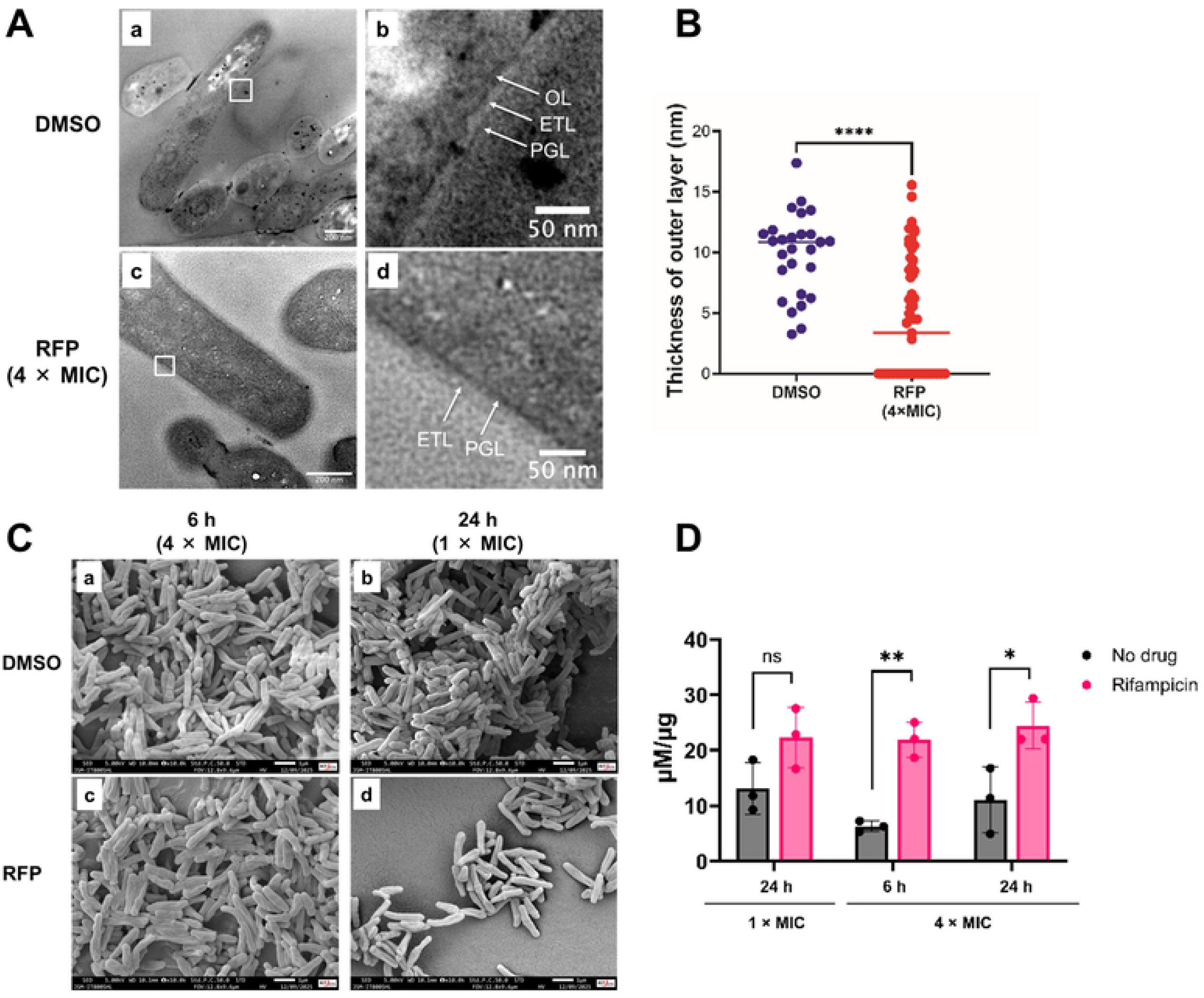
Effects of rifampicin treatment on the cell envelope ultrastructure and intracellular levels of UDP-GlcNA in *M. abscessus*. A) TEM images of *Mab* ATCC 19977 cells treated with DMSO or rifampicin. Right panels show magnified views of the white-boxed regions in the corresponding left panels. B) Quantification of outer layer thickness under each condition (DMSO or rifampicin). C) SEM images of *Mab* ATCC 19977 cells following rifampicin treatment. Left panels show cells treated with 4× MIC for 6 h, and right panels show cells treated with 1× MIC for 24 h. D) Intracellular levels of UDP-GlcNAc under each condition. Red bars indicate rifampicin-treated samples, and gray bars indicate no-drug controls. Statistical analysis was conducted in Welch’s t-test (****p < 0.0001, ** p < 0.01, *p < 0.05)

**Figure 4.**
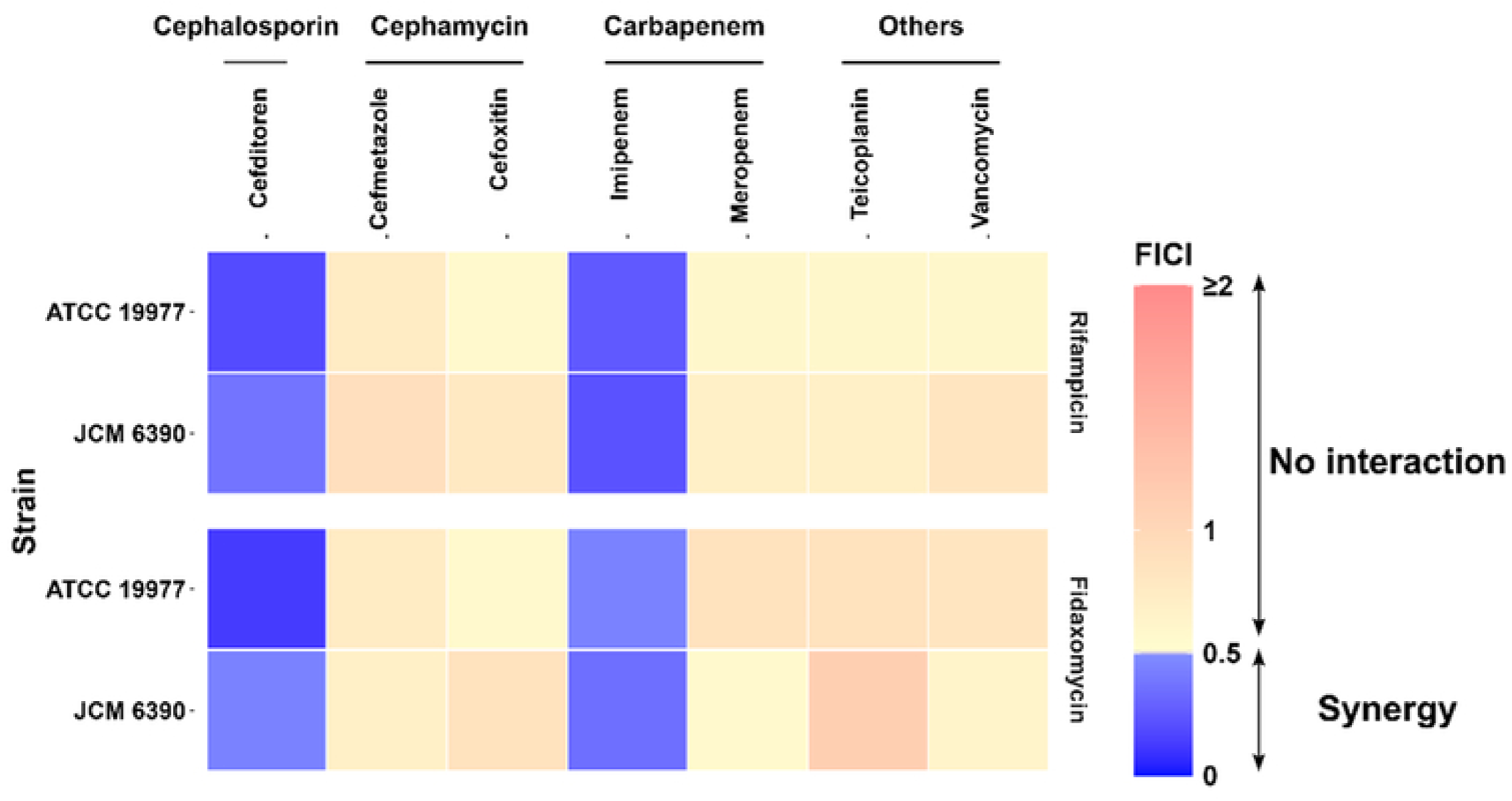
Drug interactions assessments of cell wall-targeting antimicrobial agents with rifampicin or fidaxomicin in *M. abscessus*. Upper panels show checkerboard assays performed in combination with rifampicin, while lower panels show assays performed in combination with fidaxomicin. Each panel displays FICI values, where values ≤ 0.5 (blue) indicate synergistic interactions and values > 0.5 (yellow to red) indicate no interaction.

### Rifampicin induces cell envelope ultrastructural alterations

Because inhibition of RNA polymerase increased the susceptibility of *M. abscessus* to certain β-lactams, we next examined whether rifampicin exposure induces measurable alterations in *M. abscessus* cell envelope architecture by using transmission electron microscopy (TEM) and scanning electron microscopy (SEM). TEM analysis revealed that the *M. abscessus* cell envelope displayed a characteristic three-layered architecture, consisting of an electron-dense outer layer (OL), an electron-transparent layer (ETL), and the peptidoglycan layer (PGL) (Figure 3A-a,b). In contrast, treatment with rifampicin resulted in marked ultrastructural changes, including a pronounced reduction of the OL and an apparent alteration in ETL thickness (Figure 3A-c,d, 3B and 3C). Consistent with these observations, SEM analysis revealed that *M. abscessus* cells exposed to rifampicin exhibited a highly shrunken and irregular surface topology, whereas untreated cells showed largely preserved surface structures (Figure 3D). Together, these results demonstrate that rifampicin exposure induces pronounced alterations in the cell envelope ultrastructure of *M. abscessus*.

To assess the pronounced ultrastructural changes observed upon rifampicin exposure, we quantified intracellular levels of UDP-N-acetylglucosamine (UDP-GlcNAc), a key metabolic intermediate in peptidoglycan biosynthesis, using LC-MS/MS. Compared with untreated controls, rifampicin treatment at 1× MIC for 24 hours resulted in elevated levels of UDP-GlcNAc. Moreover, exposure to rifampicin at 4× MIC for either 6 or 24 hours led to a statistically significant increase in UDP-GlcNAc abundance (Figure 3E).

## Discussion

Despite the clinical importance of rifamycins in the treatment of mycobacterial infections, their therapeutic utility against *M. abscessus* remains limited due to poorly defined intrinsic resistance mechanisms. By combining genomewide Tn-Seq with quantitative validation, we identified a high-confidence core set of 16 genes required for survival under rifamycin stress (Figure 5). This core set includes established resistance determinants such as *arr* and *helR*, but also reveals a broader physiological network that had not previously been integrated into a unified resistance framework. These findings demonstrate that intrinsic rifamycin resistance in *M. abscessus* is not governed by a single dominant mechanism but instead emerges from coordinated cellular processes. Among the identified determinants, *MAB_2807* emerged as a prominent efflux-based contributor to rifamycin resistance. Disruption of *MAB_2807* increased intracellular rifampicin accumulation and conferred broad-spectrum antibiotic susceptibility, with the most effects observed for rifamycins. The orthologs of *MAB_2807* in *M. tuberculosis* (*Rv1410c*) and *M. smegmatis* (*MSMEG_3069*) have also been shown to confer multidrug resistance, suggesting that these genes are functionally conserved across the genus (22, 23).

**Figure 5.**
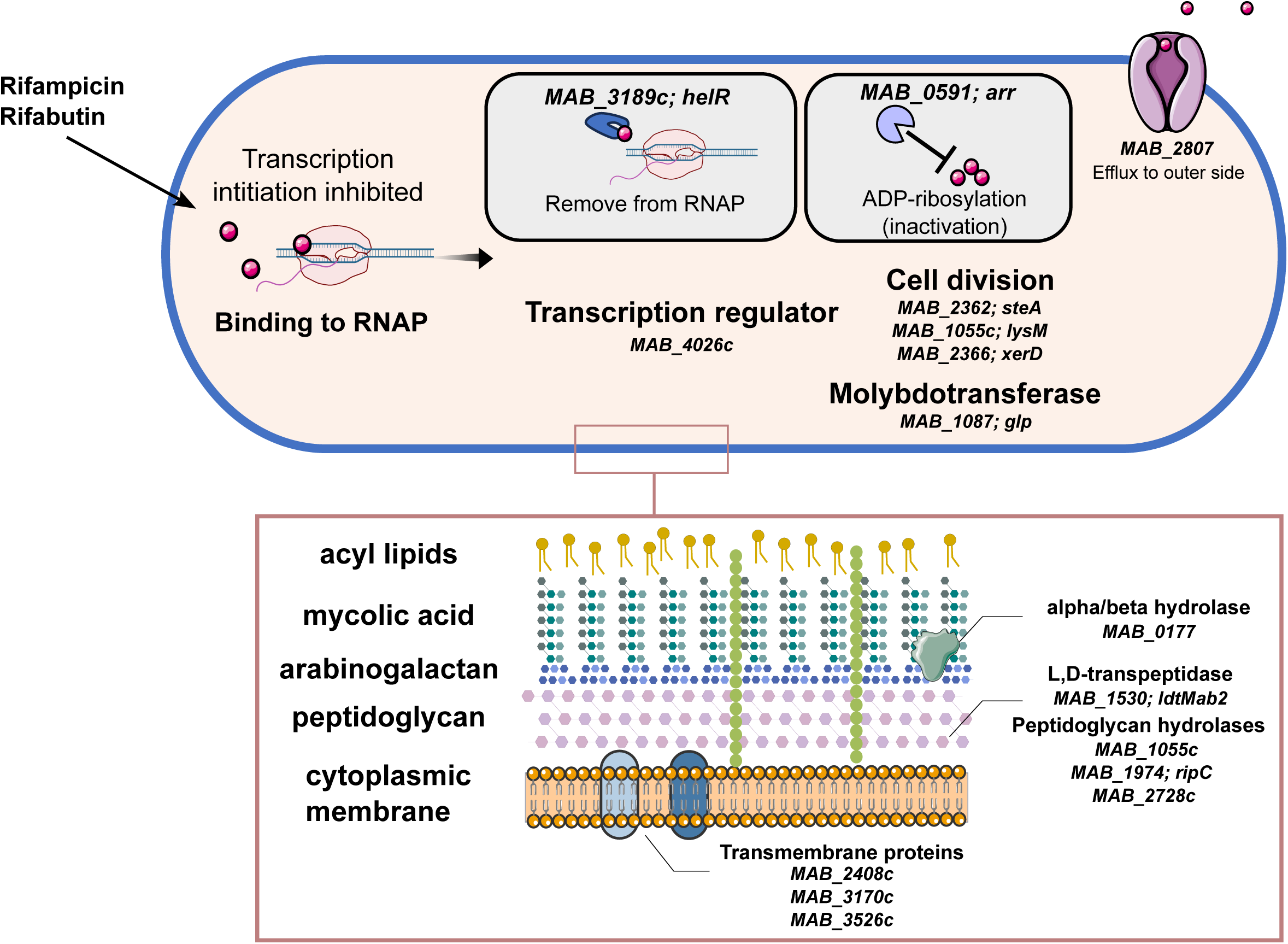
Integrated functional map of intrinsic rifamycin resistance genes. The schematic summarizes genetic determinants identified by Tn-Seq and their functional relationships to cell envelope organization, peptidoglycan metabolism, and intrinsic rifamycin resistance.

Beyond drug inactivation and efflux, the core gene set was enriched for envelope-associated determinants. Consistent with this genetic signature, we found that many cell wall targeting antimicrobial agents enhanced rifamycin activity. Unexpectedly, we found that IPM and CDTR showed synergy with rifamycin. This observation suggests that inhibition of RNA polymerase creates conditional vulnerabilities within specific peptidoglycan maintenance pathways. In mycobacteria, peptidoglycan cross-linking is dominated by noncanonical 3→3 linkages mediated by L,D-transpeptidases rather than classical 4→3 linkages mediated by D,D-transpeptidases (24–26). One of the L,D-transpeptidases (*ldtMab2*) was identified as part of the core intrinsic resistance gene set, and *ponA1, ponA2, dabB1*, which were detected only under higher rifampicin concentrations. Although selective synergy was maintained in *ΔldtMab2* (Figure S3, Table S8), this observation suggests that these enzymes likely function within a broader maintenance network that becomes increasingly important when transcriptional capacity is constrained. Consistent with this idea, rifampicin exposure induced marked ultrastructural alterations in the cell envelope and led to the accumulation of UDP-GlcNAc, a central precursor of peptidoglycan biosynthesis. Similar accumulation of UDP-GlcNAc has been reported in *rpoB* mutant strains of *Bacillus subtilis* that display collateral sensitivity to β-lactams (27), and exogenous UDP-GlcNAc has been shown to sensitize antibiotic-tolerant *E. coli* populations to β-lactam treatment (28). Although our data do not establish a direct biochemical pathway connecting RNA polymerase inhibition to specific cell wall enzymatic targets, these observations suggest that transcriptional inhibition by rifamycins perturbs envelope metabolic balance, generating a physiological state in which cell wall maintenance becomes increasingly critical.

Comparison with previously published RNA-seq datasets further clarifies the distinction between inducible and conditionally required determinants (9). Rifamycin exposure robustly induces *arr* and *helR* expression, consistent with their direct protective roles. In contrast, most Tn-Seq-defined determinants were not strongly transcriptionally upregulated, indicating that their contribution reflects physiological requirement rather than inducible activation. This separation between inducible counter-resistance factors and conditionally essential maintenance genes provides a conceptual framework for understanding intrinsic rifamycin resistance as the combined output of active drug neutralization and envelope homeostasis.

Collectively, this study defines the genomewide genetic architecture underlying intrinsic rifamycin resistance in *M. abscessus*. These findings provide a systematic foundation for rational drug combination strategies with rifamycin and inform the development and evaluation of next-generation rifamycin derivatives designed to overcome intrinsic resistance.

## Materials and Methods

### Bacterial strains, plasmids, and growth conditions

Bacterial strains and plasmids used in this study are listed in Tables S10 and S11. *M. abscessus subsp. abscessus* JCM 13569^T^ (*Mab* ATCC 19977) and JCM 6390 were obtained from the Japan Collection of Microorganisms of the Riken Bio-Resource Center (BRC-JCM; Ibaraki, Japan). Bacterial strains were cultured at 37 °C in Middlebrook 7H9 broth (BD Difco) supplemented with 10% (v/v) oleate-albumin-dextrose-catalase (OADC), 0.2% (v/v) glycerol, and 0.05% (v/v) tyloxapol. *M. smegmatis* mc^2^155, which was used for amplification of *Himar1* mycobacteriophage, was cultured in Middlebrook 7H9 broth (BD Difco) supplemented with glucose (0.2% v/v) at 30 °C. *Escherichia coli* strains were cultured in LB broth or on LB agar plate at 37 °C. When required, antimicrobial agents were added at the following final concentrations: kanamycin (50 µg/ml), anhydrotetracycline (ATc; 500 ng/ml), and zeocin (50 µg/ml) for selection of *Mab* ATCC 19977 transformants carrying pLAM12-Zeo-derived plasmids; and zeocin (25 µg/ml) for *E. coli* during plasmid construction.

### Construction of saturated transposon insertion mutant libraries of *Mab* ATCC 19977

Saturated transposon insertion mutant libraries were generated in *Mab* ATCC 19977 using the mycobacteriophage phAE180 carrying a *Himar1*-derived mariner transposon as described previously (32, 33). Phage stocks (8-10 × 10¹⁰ PFU/ml) were used to transduce exponentially growing cultures (OD₆₀₀ = 0.8-1.7). Following transduction, cells were resuspended in 7H9 medium and plated onto 7H10 agar supplemented with 0.05% tyloxapol and kanamycin (50 µg/ml). Plates were incubated at 37 °C for 4-6 days. Colonies were scraped, pooled, and stored at −80 °C until use. Two independent biological libraries were generated.

### Antibiotic selection of transposon insertion mutant libraries

The growth-inhibitory effects of rifampicin and rifabutin on *Mab* ATCC 19977 when grown on 7H9 agar plates were assessed as described below. Cultures were grown to mid-log phase and serially diluted starting from an initial OD₆₀₀ of 1.0. Five microliters of each dilution were spotted onto 7H9 agar plates containing the indicated concentrations of antibiotics. Plates were incubated at 37 °C for 4 days.

Frozen stock of the *Mab* ATCC 19977 transposon insertion mutant library was thawed and diluted to 1.0 × 10^6^ colony forming unit (CFU)/ml in 5 ml fresh 7H9 medium. The diluted libraries were plated on 7H9 agar plates supplemented with kanamycin and 0.5 µg/ml, 5 µg/ml rifampicin, or 0.5 µg/ml rifabutin. All agar plates were incubated at 37 °C for 7 days (5 µg/ml rifampicin) or 4 days (0.5 µg/ml rifampicin or 0.5 µg/ml rifabutin). All colonies grown on each plate were suspended in fresh 7H9 medium, aliquoted into 2 ml cryo-tubes, and stored at -80 °C for further genomic DNA extraction.

### Tn-Seq

Genomic DNA was extracted using a phenol-chloroform extraction method following mechanical disruption by bead beating (0.1-mm zirconia beads; Vortex Mixer GENIE2 with Microtube Attachment, maximum speed for 7 minutes), as previously described (33, 34). Tn-Seq libraries were prepared as described previously (33, 35), enriching transposon-genome junction fragments by PCR amplification. Libraries were sequenced on an Illumina NextSeq 2000 platform using 125-bp paired-end reads (NextSeq 1000/2000 P1 XLEAP-SBS reagents). Raw sequencing reads were processed using the TPP module of the TRANSIT v3.3.4 software package. Transposon insertion sites were identified by mapping reads to the *Mab* ATCC 19977 reference genome using the Burrows-Wheeler Aligner (BWA) (36). Gene essentiality under non-selective conditions was determined using the Hidden Markov Model (HMM) implemented in TRANSIT, which classifies genes based on insertion density and local read count distributions. HMM confidence scores were calculated to evaluate the robustness of each classification.

Conditional essentiality under antibiotic treated conditions was assessed using the resampling method implemented in TRANSIT. *P*-values were corrected for multiple hypothesis testing using the Benjamini-Hochberg procedure with a false discovery rate (FDR) threshold of 5%. Genes with adjusted *p*-values (Padj) < 0.05 were considered significantly different in fitness relative to the untreated control. Mutual orthologues between *Mab* ATCC 19977 and *M. tuberculosis* H37Rv were defined by reciprocal best-hit analysis using BLASTp with an e-value cutoff of 1×10⁻¹⁰.

### Isolation of rifampicin-susceptible transposon insertion mutants

To isolate individual transposon insertion mutants with increased rifampicin susceptibility, a total of 702 *Mab* ATCC 19977 transposon insertion mutant colonies were screened by replica patching onto 7H9 agar plates containing kanamycin alone or kanamycin supplemented with rifampicin (2 µg/ml). Mutants that exhibited impaired growth on kanamycin plus rifampicin plates were selected for secondary screening. For secondary screening, rifampicin susceptibility was assessed by MIC determination. The *Himar1* transposon insertion site was identified as previously described (37). Briefly, genomic DNA from each mutant was digested with *Bss*HII and self-ligated to generate circular DNA fragments. Circularized fragments containing the *ori6K* origin within the *Himar1* transposon were transformed into *E. coli* DH5αλpir. Plasmids were purified from the resulting transformants, and genomic regions adjacent to the 3′ end of the transposon were sequenced using the primer KanSeq_Rev (38). Insertion sites were determined by aligning Sanger sequencing reads to the *Mab* ATCC 19977 reference genome (GenBank accession number NC_010397.1). The genomic locations of the confirmed insertion sites were summarized in Figure S6.

### Construction of gene deletion mutants using a modified ORBIT system

*Mab* ATCC 19977 gene deletion mutants of *arr*, *ldtMab2*, *ripC*, and *helR* were constructed using an ORBIT system (20). Due to the intrinsic tetracycline resistance of *M. abscessus*, the original tetracycline-inducible ORBIT system was ineffective, likely because of insufficient promoter induction. Thus, we constructed an acetamide-inducible RecT annealase and phage Bxb1 integrase expression plasmid; a DNA fragment encoding both enzymes was amplified from pKM461 using primers P1558 and P1559. The fragment was inserted into an *Nde*I-digested pLAM12 (Addgene plasmid #26908) using the In-Fusion Snap Assembly Master Mix (Takara Bio), generating plasmid pEPU196. The strain harboring pEPU196 was cultured in 10 ml of 7H9 medium supplemented with kanamycin (50 µg/ml) at 37 °C until mid-exponential phase. Acetamide was then added to a final concentration of 0.2% (v/v), and cultures were incubated for an additional 3 hours to induce expression of RecT annealase and Bxb1 integrase. Cultures were subsequently transferred to conical tubes and incubated on ice for 90 minutes. Cells were harvested by centrifugation at 3,080 g for 10 minutes at 4 °C. The supernatant was completely removed, and cells were washed three times with pre-chilled 10% glycerol. The wash volume was reduced by approximately half at each successive step. After the final wash, cells were resuspended in 100 µl of 10% glycerol to generate electrocompetent cells. For electroporation, 100 µl of competent cells were mixed with 1 µg of targeting oligonucleotide and 200 ng of the payload plasmid pKM496 and transferred to ice-cooled 0.2-cm gap cuvettes. Electroporation was performed at 2,500 V, 25 µF, and 1,000 Ω. Cells were recovered in 7H9 broth overnight at 37 °C and plated on 7H9 agar supplemented with zeocin (25 µg/ml). Plates were incubated at 37 °C for 4-5 days. Recombinant clones were verified by colony PCR using GoTaq Colorless Master Mix (Promega) supplemented with 5% (v/v) DMSO. PCR conditions were as follows: 95 °C for 10 minutes; 20 cycles of 95 °C for 30 seconds, 58 °C for 30 seconds, and 72 °C for 5 minutes; followed by a final extension at 72 °C for 5 minutes. Correct integration was confirmed by detection of appropriately sized PCR products spanning both chromosomal-plasmid junctions. Primer and oligo sequences are listed in Table S7.

### Construction of *MAB_2807* complemented strain

Complementation of the *MAB_2807*::Tn mutant was performed by introducing a replicative mycobacterial expression plasmid carrying the full-length *MAB_2807* coding sequence. The gene was expressed under the control of an acetamide-inducible promoter. To generate the complementation construct, the *MAB_2807* open reading frame was amplified from *Mab* ATCC 19977 genomic DNA using gene-specific primers (Table S7). The PCR product was cloned into the mycobacterial shuttle vector pLAM12-Zeo, which carries a zeocin resistance cassette for selection. The resulting plasmid was verified by Sanger sequencing and electroporated into the *MAB_2807*::Tn mutant strain. Transformants were selected on 7H10 agar supplemented with zeocin.

### Rifampicin accumulation assay

Intracellular rifampicin accumulation was quantified in *Mab* ATCC 19977 and the *MAB_2807*::Tn mutant. Cultures were grown in 25 ml of 7H9 medium at 37 °C with shaking to mid-exponential phase. Cells were harvested by centrifugation at 2,500 g for 10 min at room temperature and resuspended in 1 ml of fresh 7H9 medium. Rifampicin was added to a final concentration of 400 µg/ml, and samples were incubated at 37 °C with shaking for 60 minutes. Following incubation, cells were washed three times with hypertonic buffer (10 mM Tris-HCl [pH7.3], 0.5 mM MgCl₂, 150 mM NaCl) to remove extracellular rifampicin and resuspended in 500 µl of the same buffer. Cells were lysed by boiling for 30 min and centrifuged at 10,000 g for 2 min to remove cellular debris. Supernatants were collected and diluted 1:2 with hypertonic buffer, and rifampicin levels were quantified by measuring absorbance at 470 nm (OD₄₇₀). The linear detection range was confirmed using a rifampicin standard curve. Rifampicin accumulation was normalized to cell counts and expressed as A₄₇₀ per 10⁵ cells. Data shown represent biological duplicates, and the experiment was independently repeated with comparable results.

### Generation of CRISPRi mutant

An *rpoB* (*MAB_3869c*) knockdown strain was generated using the CRISPR interference plasmid pLJR962 (Addgene plasmids #115162) (39, 40). Briefly, a single-guide RNA (sgRNA) targeting *rpoB* was designed using the Pebble sgRNA Design Tool (Rock Lab, Rockefeller University; https://pebble.rockefeller.edu/tools/sgrna-design/). The corresponding sgRNA oligonucleotides (Table S7) were annealed and ligated into pLJR962 according to established protocols. Correct plasmid construction was confirmed by Sanger sequencing using the primer 1834 (Table S7). The recombinant plasmid was first transformed into *E. coli* TOP10 for propagation and plasmid amplification. Purified plasmid DNA was subsequently electroporated into *Mab* ATCC 19977 with 2,500 V, 25 µF, and 1,000 Ω using ice-cooled 0.2-cm gap cuvettes. Cells were recovered in 7H9 broth overnight at 37 °C and plated on 7H9 agar supplemented with kanamycin (50 µg/ml). Plates were incubated at 37 °C for 4-5 days. Primer and oligo sequences are listed in Table S7.

### MIC and growth fitness determinations

*M. abscessus* strains were grown to mid-log phase and diluted to an OD₆₀₀ of 0.001 in 7H9 medium. In case of *rpoB* knockdown, the inhibition was induced with 500 ng/µl anhydrotetracycline (ATc). Two hundred microliters of the bacterial suspension were dispensed into 96-well flat-bottom microplates (TPP) containing twofold serial dilutions of the indicated antibiotics. Antimicrobial dilution series were prepared using a digital dispenser (Multidrop Pico 8; Thermo Fisher Scientific). Plates were incubated at 37 °C, and OD₆₀₀ was measured using a microplate reader (Synergy HTX, BioTek). For MIC determination, plates were incubated for 5 days. After incubation, plates were briefly mixed using a microplate shaker, and OD₆₀₀ was measured. MIC values were defined as the lowest antibiotic concentration that inhibited ≥90% of bacterial growth relative to the drug-free control. Each strain was tested with technical replicates per plate and independently repeated two times.

For growth fitness analysis, OD₆₀₀ measurements were recorded every 30 minutes for 96 hours. Growth curves were used to calculate the area under the curve (AUC) for each condition. Relative fitness was defined as the ratio of AUC in the presence of drug to that in the absence of drug:

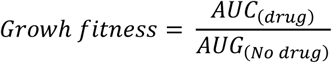

Reported values represent the mean of technical replicates, and experiments were independently repeated two times.

### SEM and TEM sample preparation and image capturing

*Mab* ATCC 19977 cultures were grown to mid-log-phase and adjusted to an OD₆₀₀ of 0.3 in 30 ml of 7H9 medium. Rifampicin was added at 1× MIC or 4× MIC, and an equivalent volume of DMSO was added to control cultures. After the indicated treatment period, cells were harvested by centrifugation at 3,080 g for 10 minutes at 4 °C and washed three times with 10 ml of phosphate buffered saline (pH 7.4). Cells were fixed in 2.5% (v/v) glutaraldehyde in 0.1 M phosphate buffer (pH 7.4). For TEM, fixed cells were washed three times with 0.1 M phosphate buffer and post-fixed in 1% (w/v) osmium tetroxide (OsO₄) at 4 °C for one hour. Samples were dehydrated through a graded ethanol series (50%, 70%, 80%, 90%, and 100%). Resin substitution was performed using a 1:1 mixture of ethanol and Spurr’s resin, followed by pure Spurr’s resin. Samples were embedded and polymerized at 70°C for 72 hours. Ultrathin sections (80 nm) were prepared by using ARTOS 3D ultramicrotome (Leica Microsystems GmbH, Wetzlar, Germany) equipped with ultra 45° diamond knife (DiATOME AG, Nidau, Switzerland), stained with Uranyless (Delta Microscopies, Mauressac, France) and Reynolds’ lead citrate, and observed using a JEM-2100Plus transmission electron microscope (JEOL, Tokyo, Japan) at an accelarating voltage of 200 kV. Cell envelope thickness was measured manually using Fiji software analyzed 100 cells in each condition (41). For SEM, dehydration was performed as described for TEM. Samples were then subjected to *tert*-butyl alcohol (TBA) substitution, which was conducted by ethanol/TBA [1:1] twice, followed by 100% TBA twice. After freeze-drying overnight, specimens were sputter-coated with gold at 10-nm thickness using Q150TS Plus sputter coater (Quorum Technologies, East Sussex, UK) and observed using a JEM-IT800SHL scanning electron microscope (JEOL) at an accelartion voltage of 5 kV. Both TEM and SEM observations were conducted in technical replicates.

### Checkerboard assay

Drug interaction assays were performed in 384-well microplates using a 7 × 7 two-dimensional concentration matrix for high-throughput screening. For selected combinations (rifampicin [RFP] or rifabutin [RBT] with imipenem [IPM] or cefditoren [CDTR]), an expanded 11 × 11 concentration matrix was used to evaluate a broader concentration range. Antibiotics were dispensed using a digital dispenser (Multidrop Pico 8; Thermo Fisher Scientific). Mid-log-phase cultures were diluted to an OD₆₀₀ of 0.001 and added to wells containing twofold serial dilutions of each drug combination.

Plates were incubated at 37 °C for 5 days, briefly mixed using a plate shaker, and bacterial growth was quantified by measuring OD₆₀₀ with a Synergy HTX microplate reader (BioTek). The fractional inhibitory concentration index (FICI) was calculated as:

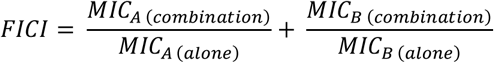

Drug interactions were classified as follows: FICI ≤ 0.5, synergy; 0.5 < FICI ≤ 2, additive (no interaction); FICI > 2, antagonism.

### Metabolite extraction and quantification of UDP-GlcNAc

*Mab* ATCC 19977 cultures were grown to mid-log phase (OD₆₀₀, 0.3-0.5) at 37 °C. Rifampicin was added at 1× MIC (32 µg/ml) or 4× MIC (128 µg/ml), and an equivalent volume of DMSO was added to control cultures in 30 ml of 7H9 medium. After the indicated treatment period, cells were harvested by centrifugation at 20,600 g for 5 minutes at 4 °C and washed once with ice-cold phosphate buffered saline (pH7.4). Cell pellets were resuspended in 1 ml of pre-cooled extraction solvent (acetonitrile:methanol:water, 40:40:20, v/v/v). To extract lysates, bead beating was conducted using 0.1 mm zirconia beads and Vortex Mixer GENIE2 with Microtube Attachment at max speed for 1 minute, followed by 2 minutes on ice, repeated four times. Lysates were centrifuged at 15,500 g for 10 minutes at 4 °C, and the supernatants were collected for subsequent LC-MS/MS analysis of UDP-GlcNAc levels.

### Liquid chromatography and mass spectrometry

One microliter of extracted metabolites was injected into an LCMS-8060NX system (SHIMADZU) for chromatographic separation and mass spectrometric detection. Aquity UPLC BEH Amide (2.1 x 100 mm, Waters) and a 20:80 mixture of solvent A (100mM ammonium formate aqueous solution) and solvent B (acetonitrile containing 0.1% formic acid) were used as the analytical column and the mobile phase, respectively. The gradient was changed from -80% B to 10% B in 5 minutes The flow rate was maintained at 0.3 mL/min. UDP-GlcNAc and ADP-Glc (internal standard) were analyzed in negative-ion, multiple-reaction-monitoring (MRM) mode. The mass transitions used were *m/z* 606 > 385 for UDP-GlcNac and *m/z* 588 > 346 for ADP-Glc. Mass axis calibration was performed using continuous reference mass infusion according to the manufacturer’s instructions. Ion abundance corresponding to UDP-GlcNAc was quantified using LobSolutions version 5.135 (SHIMADZU). UDP-GlcNAc levels were normalized to total protein concentration determined using the Pierce™ BCA Protein Assay Kit (Thermo Fisher Scientific). Statistical analyses were performed using Welch’s unpaired t-test with at least three independent biological replicates. Differences were considered statistically significant at *p*-value < 0.05.

### Statistical analysis

Statistical analyses and graphical representations were performed using GraphPad Prism (version 10.6.0), Inkscape 1.3.2 (091e20e, 2023-11-25, custom) with additional analyses conducted in R software.

## Data availability

All sequence data are available in the NCBI BioProject database under accession number PRJDB40469 with individual Sequence Read Archive (SRA) dataset accession numbers DRR959860-DRR959867.

## Acknowledgements

*M. smegmatis* mc^2^155 and mycobacteriophage phAE180 were gifts from William R. Jacobs, Jr., of the Albert Einstein College of Medicine.

